# A Data-Independent-Acquisition-based proteomic approach towards understanding the acclimation strategy of *Microchloropsis gaditana* CCMP526 in hypersaline conditions

**DOI:** 10.1101/2020.03.18.996223

**Authors:** Anbarasu Karthikaichamy, John Beardall, Ross Coppel, Santosh Noronha, Dieter Bulach, Ralf B. Schittenhelm, Sanjeeva Srivastava

## Abstract

Salinity is one of the significant factors that affect growth and cellular metabolism, including photosynthesis and lipid accumulation, in microalgae and higher plants. *Microchloropsis gaditana* CCMP526 can acclimatize to different salinity levels by accumulating compatible solutes, carbohydrates, and lipid as an energy storage molecule. We used proteomics to understand the molecular basis for acclimation of *M. gaditana* to increased salinity levels (55 and 100 PSU (Practical Salinity Unit). Correspondence analysis (CA) was used for identification of salinity-responsive proteins (SRPs). The highest number of altered proteins was observed in 100 PSU. Gene Ontology (GO) enrichment analysis revealed a separate path of acclimation for cells exposed to 55 and 100 PSU. Osmolyte and lipid biosynthesis was up-regulated in high saline conditions. However, concomitantly lipid oxidation pathways were also up-regulated at high saline conditions, providing acetyl-CoA for energy metabolism through the TCA cycle. Carbon fixation and photosynthesis were tightly regulated, while chlorophyll biosynthesis was affected under high salinity conditions. Importantly, temporal proteome analysis of salinity-challenged *M. gaditana* revealed vital salinity-responsive proteins which could be used for strain engineering for improved salinity resistance.

## Introduction

Biofuel sourced from algae is considered as an effective renewable alternative to fossil fuels (Georgianna & Mayfield, 2012). Microalgae can grow in a variety of environmental conditions all year-round and produce valuable products such as biofuel, animal feeds, and pharmaceuticals (Brennan & Owende, 2010; Doan et al., 2011). Moreover, many species can accumulate high amounts of lipid under stress, and this phenomenon has been extensively researched in promising biofuel candidates under various stress conditions (Corteggiani Carpinelli et al., 2014; Huertas et al., 2000; Simionato et al., 2011). However, limitations such as low lipid productivity and high costs associated with downstream processing have hindered the commercialization of algae-based biofuel (Wijffels & Barbosa, 2010). A simple approach for enhancing productivity coupled with high growth rate would be to manipulate the environmental parameters. Salinity is one of the primary growth influencing factors for marine microalgae, which could be controlled in an open race-way pond (Kim et al., 2016; Perrineau et al., 2014).

Extensive research has been done on increasing lipid accumulation in various microalgal species using salinity stress (Kim et al., 2016; Perrineau et al., 2014). Notably, the halotolerant alga *Dunaliella* sp. is known to perform well in high saline conditions. It accumulates a high amount (~5 M) of glycerol when exposed to hypersaline conditions (Takagi & Yoshida, 2006). Lipid accumulation was enhanced to 59.4% in *Chlamydomonas* sp. JSC4 by gradually increasing the salinity levels up to 2% (w/v) of sea salt (Ho et al., 2017). Accumulation of lipid or glycerol in such conditions involves a major metabolic shift in re-directing the carbon flux, which causes alteration in the abundance of proteins/enzymes involved in the metabolism of lipid or glycerol.

Appropriately, various proteomic studies can be performed to understand the acclimatization process of microalgae in high saline conditions. The proteome of the halotolerant alga *Dunaliella* sp. has been studied to understand the molecular basis behind tolerance to high saline conditions (Katz et al., 2007; Wei et al., 2017). Sithtisarn et al. (2017) performed comparative proteome analysis of a control, and a salinity tolerant strain of *Chlamydomonas reinhardtii* to understand the salinity tolerance mechanism involved. Alterations in proteome of terrestrial plants have also been extensively studied (Aghaei et al., 2008; Silveira & Carvalho, 2016). The response to salinity appears similar in both microalgae and higher plants. Several biosynthetic pathways were affected under high salinity conditions, including photosynthesis, lipid, carbohydrate, and amino acid metabolism (Pandit et al., 2017).

*Microchloropsis gaditana* CCMP526, previously known as *Nannochloropsis gaditana* CCMP526, is one of the six algal species from the genus *Nannochloropsis* (Fawley et al., 2015). It is a marine microalga known for high oil accumulating capability and especially its eicosapentaenoic acid (EPA) content. Physiological alterations in the lipid and carbohydrate accumulation have been observed in high salinity conditions, which prompted us to investigate the genes and proteins involved in the acclimatization mechanism. A systematic investigation using NGS-based transcriptomic (RNA-Seq) approach was performed to identify the molecular players in salinity tolerance. The results showed active shunting of carbon flux towards lipid accumulation and osmolyte biosynthesis. In this work, we aim to identify the salinity-responsive proteins (SRPs) using a DIA (data-independent acquisition)-proteomic approach. The traditional proteomic analysis uses data-dependent acquisition (DDA), which is often limited by the number of proteins and peptides that can be analyzed in one run. In DIA analysis, all peptides within a defined *m/z* range are fragmented, resulting in an increase in the number quantifiable peptides and a more accurate peptide quantification (Doerr, 2014).

Temporal proteomic analysis was performed on *M. gaditana* subjected to different salinity levels (38, 55 and 100 PSU). Time points and salinity levels for sampling were chosen based on the lipid and carbohydrate accumulation from our previous physiological analysis. Systematic analysis of the temporal proteome data revealed vital salinity-responsive proteins (SRPs) and the dynamics of essential proteins involved in the cellular metabolic shift towards acclimatization.

## Materials and methods

### Cultivation and harvesting

*M. gaditana* was cultivated in 0.2 μm-filtered sea water (collected from the Gippsland Lakes, Gippsland, Victoria, Australia), supplemented with Guillard's f/2 nutrients ^1^ and 17 mM sodium nitrate, at 25°C using 500 ml glass bottles (Schott Duran, Germany). Cultures (300 ml) were mixed by bubbling with sterile air (0.2 μm filtered) supplied at a flow rate of 2.5 L min^−1^. Illumination was provided at 150 μmol · photons · m^−2^ · s^−1^ (Philips, TLD36W, Amsterdam, The Netherlands) with a light/dark cycle of 12/12 h. Sodium chloride was added to the existing f/2 media (control, 38 PSU) to make f/2 media of different salinities (55 and 100 PSU). A portable refractometer (RHS-10ATC) was used to assess the salinity of the culture media. Growth curve experiment (cell count and OD_685_) was performed in standard conditions (38 PSU, 25°C, and 150 μmol · photons · m^−2^ · s^−1^) to determine the growth phases in *M. gaditana* (data not shown). Cells at mid exponential phase were inoculated into f/2 media of different salinities at a concentration of approximately 4×10^6^ cells/ml. Cells were then sampled from three independent cultures at specific time points (0, 1, 6, 24 and 72 h) that were chosen from our earlier physiological study ^2^. The samples were centrifuged (4000 ×g, 4°C, 5 min) in an Heraeus Multifuge model 3SR Plus (thermos Scientific, Australia) and the resulting cell pellet was washed thrice with sterile distilled water to remove any residual salt from the growth medium. The pellet was then stored at −80°C until further processing.

### Protein isolation and Sample preparation for mass spectrometry

Protein was isolated using a Bioline II DNA/RNA/Protein extraction kit (Bioline, Australia) following the manufacturer’s protocol. The isolated protein was quantified using the absorbance at 280 nm (Thermo Scientific, Australia). The samples were stored in −80°C until further processing. Approximately 50 mg of protein was denatured using TCEP (Thermo Scientific, Cat. #77720) to a final concentration of 10 mM. The solution was then incubated at 50-65°C for 30 min. Chloroacetamide (CAA) was added to a final concentration of 40 mM to alkylate the reduced protein. The mixture was incubated at room temperature in the dark for 20 min. Trypsin at a dilution of 1:100 was added and incubated overnight at 37°C with shaking. The reaction was stopped by adding 1% formic acid (FA) (Wei et al.). The samples were then desalted using Ziptips (Agilent, OMIX-Mini Bed 96 C18, A57003MBK). The residual acetonitrile (ACN) was removed using a vacuum concentrator. The sample was dissolved into 20 ml of 0.1% FA and sonicated in a water bath for 10 min. Any insoluble precipitates were removed by centrifuging for 5 min. iRT (indexed retention time) peptides were added before transferring the sample into an MS vial.

### Mass spectrometry Data Acquisition

#### Data-dependent acquisition (DDA) mass spectrometry

Using a Dionex UltiMate 3000 RSLCnano system equipped with a Dionex UltiMate 3000 RS autosampler, the samples were loaded via an Acclaim PepMap 100 trap column (100 μm × 2 cm, nanoViper, C18, 5 μm, 100å; Thermo Scientific) onto an Acclaim PepMap RSLC analytical column (75 μm × 50 cm, nanoViper, C18, 2 μm, 100å; Thermo Scientific). The peptides were separated by elution with increasing concentrations of 80% ACN / 0.1% FA at a flow rate of 250 nl/min for 158 min and analyzed with an Orbitrap Fusion Tribrid mass spectrometer (Thermo Scientific). Each cycle was set to a fixed cycle time of 4 sec consisting of an Orbitrap full ms1 scan (resolution: 120.000; AGC target: 1e6; maximum IT: 54 ms; scan range: 375-1575 m/z) followed by several Orbitrap ms2 scans (resolution: 30.000; AGC target: 4e5; maximum IT: 118 ms; isolation window: 1.4 m/z; HCD Collision Energy: 32%). To minimize repeated sequencing of the peptides, dynamic exclusion was set to 15 sec, and the ‘exclude isotopes’ option was activated. The raw and analyzed DDA data files have been deposited to the ProteomeXchange Consortium (http://proteomecentral.proteomexchange.org) via the PRIDE partner repository (Vizcaíno et al., 2016)^3^ with the dataset identifier PXD017164 (Username, Password: 1MhuJS1T).

#### Quantification of proteins using data-independent acquisition (DIA) mass spectrometry

The identical instrument setup as described above (Dionex UltiMate 3000 LC system coupled to an Orbitrap Fusion Tribrid mass spectrometer) was used to quantify proteins using data-independent acquisition (DIA). 25 sequential DIA windows with an isolation width of 24 m/z between 375 - 975 m/z were acquired (resolution: 30.000; AGC target: 1e6; maximum IT: 54 ms; HCD Collision energy: 32%; scan range: 200-2000 m/z) following a full ms1 scan (resolution: 120.000; AGC target: 1e6; maximum IT: 54 ms; scan range: 375-1575 m/z). A 158 min gradient of increasing concentrations of 80% ACN / 0.1% FA was used to separate the peptides for the DIA acquisition. The raw and analyzed DIA data files have been deposited to the ProteomeXchange Consortium (http://proteomecentral.proteomexchange.org) via the PRIDE partner repository (Vizcaíno et al., 2016) with the dataset identifier PXD017297 (Username, Password: Yszk3q3x).

#### Mass spectrometric data analysis

Acquired DDA .raw files were searched against the *N. gaditana* UniProtKB/SwissProt database (v2017_07) using Byonic (Protein Metrics) embedded into Proteome Discoverer (Thermo Scientific) to obtain peptide sequence information. Only peptides identified at a false discovery rate (FDR) of 1% based on a decoy database were considered for further analysis. Spectronaut Orion (Biognosys) was used to create the corresponding spectral library as well as to evaluate all DIA data using in-house parameters. To correct for differences in sample density and loading, the peak areas for each peptide were normalized based on the assumption that on average, a similar number of peptides are up- and down-regulated. Two-sided t-tests were used to calculate p-values and the FDR for each time point (0, 1, 6, 24 and 72 h) and salinity level (55 and 100 PSU) against control (38 PSU), based on multiple hypotheses testing corrections by Benjamini-Hochberg method (implemented in R). GO terms were enriched using a Fisher exact test with p-value <0.05.

## Results and discussions

### General description of the proteomics results

DIA based proteomic analysis was employed to investigate the temporal acclimatization strategy of *M. gadit*ana at different salinity levels (55 and 100 PSU) compared to controls at 38 PSU, and at various time points (0, 1, 6, 24 and 72 h). These time points represent the critical stages of change in cellular state in regard to alterations in photosynthetic rate, carbohydrate, and lipid accumulation (Karthikaichamy et al., 2018). Changes in protein level are represented as average log_2_(fold change) compared with the control (38 PSU) sample at corresponding time points (figure 1A). On average, the proteins were up-regulated in cells exposed to 100 PSU (except at 0 and 24 h). A similar trend was observed in cells in 55 PSU, except at 6h, where the average log_2_(fold change) value indicated significant down-regulation of proteins (p-value=0.000002). Noticeably, the average log_2_(fold change) in 100 PSU was significantly higher than 55 PSU at 72 h (p-value <0.000001). This sudden up-regulation of protein during the later phase of growth in 100 PSU indicates a strong response towards acclimation. To identify the proteins that were expressed in all timepoints, Venn diagram was constructed (figure 1B). A total of 1874 proteins were expressed across all the time points, and these proteins were selected for temporal analysis.

**Figure 1.**
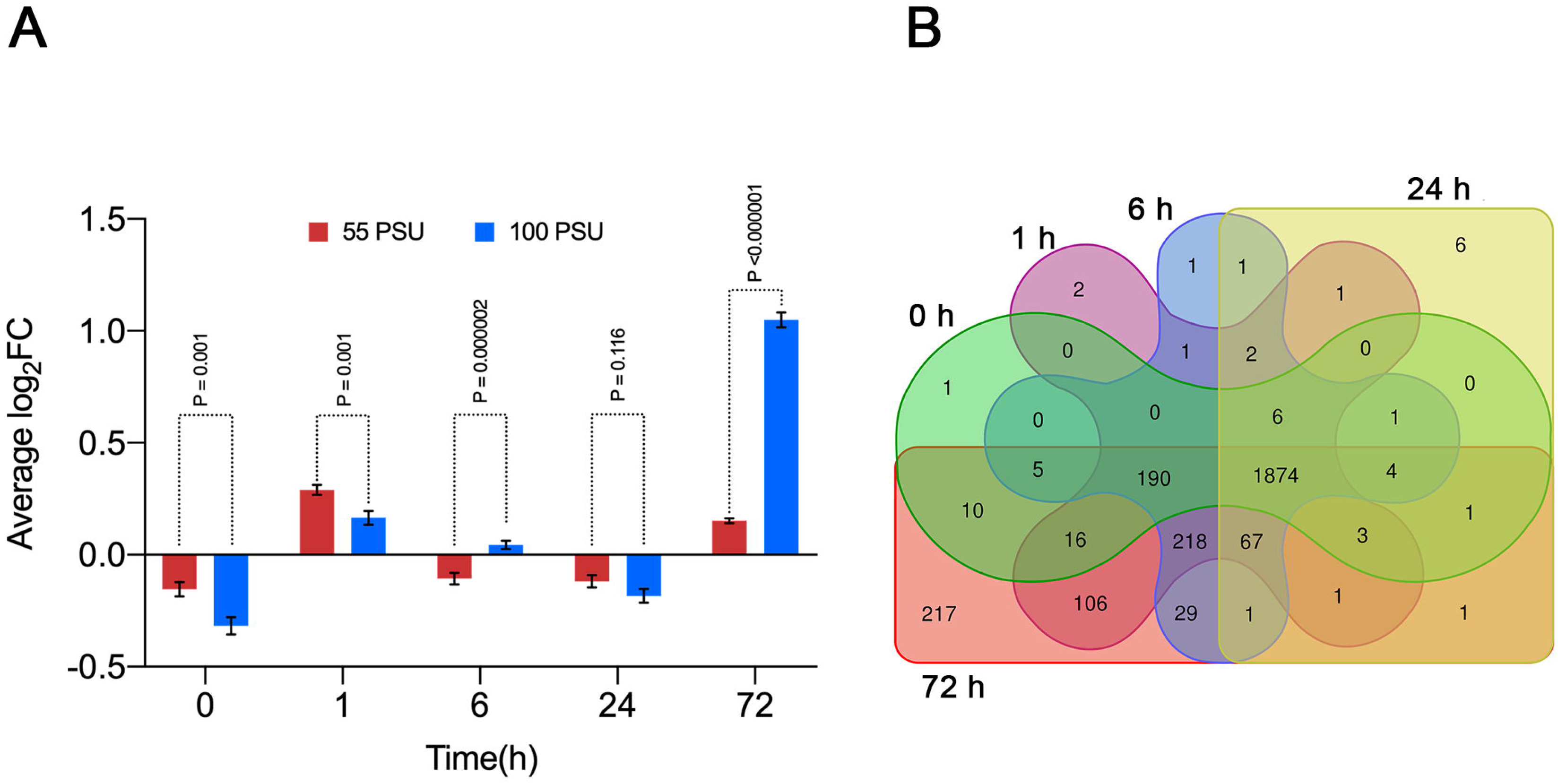
**A.** Differential protein expression at different time points and salinity levels (55 and 100 PSU) compared to control (38 PSU). Values plotted are the average expression at a particular time point, and error bars indicate SEM; **B.** Venn diagram showing the overlapping differentially abundant proteins in different time points.

The proteins that satisfied the cut-off criteria (p-value<0.05 and absolute log_2_(fold change) >1) in at-least single time-point were selected for further analysis. *M. gaditana* grown in 55 PSU had 456 significantly expressed proteins compared to 1359 proteins in cells exposed to 100 PSU relative to control condition (38 PSU). The highest number of proteins that were differentially expressed were in cells grown under 100 PSU at 72 h (264 down-regulated and 1110 up-regulated proteins), suggesting a major proteomic shift towards acclimation in high saline conditions. There was also a robust initial response to change in salinity. Ideally the protein expression levels should remain the same at 0h, irrespective of the salinity. However, we observed differential expression of proteins at 0h in 55 and 100 PSU. This is explained by the time required for harvesting and processing the cells for proteomics experiment. In 55 and 100 PSU, 157 and 454 proteins were down-regulated respectively (compared to controls) at 0 h, which was higher than at any of the other time points (Figure 2). Moreover, 76 and 120 proteins were statistically up-regulated in cells exposed to 55 and 100 PSU respectively at 0 h, indicating the initial acclimation strategy towards high salinity conditions. List of differentially expressed proteins in *M. gaditana* in high salinity conditions (55 and 100 PSU) are given in supplementary table S1.

**Figure 2.**
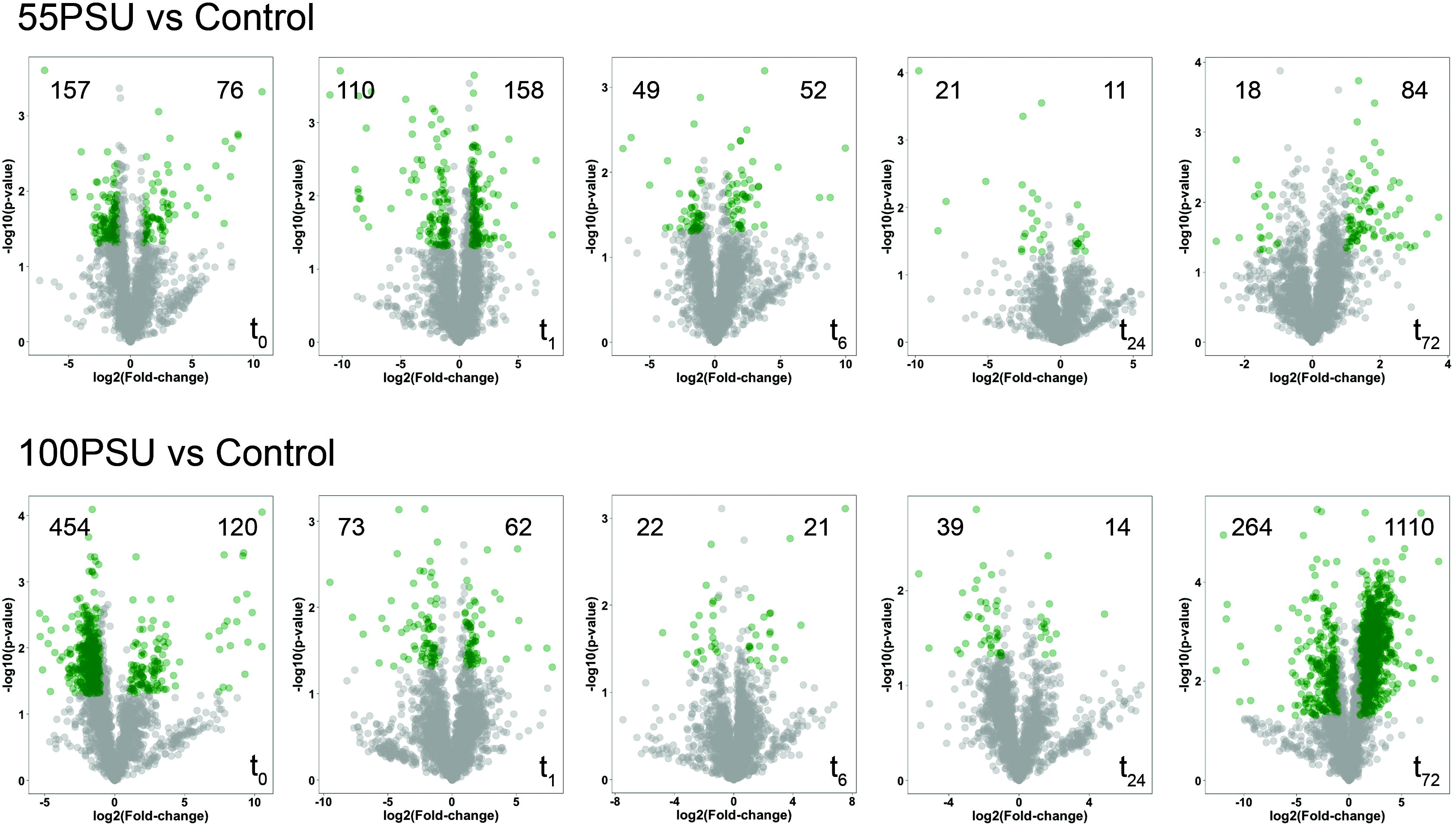
Volcano plot showing significantly regulated proteins as green dots (p-value<0.05 and absolute log2(fold change) >1). Values on either side of the plot represent the number of significantly up- and down-regulated proteins.

### GO-term enrichment reveals the path towards acclimation

The disparity in the number of differentially expressed proteins at various time points and different salinity levels suggests that *M. gaditana* employs a different strategy for acclimation to salinities of 55 and 100 PSU. To understand the path towards acclimation, statistically significant proteins at different time points were GO-enriched in Blast2GO using a filtering cut-off of p-value<0.05 (Conesa et al., 2005). Interactive maps were created with the significantly enriched biological process (BP) GO-terms (supplementary tables S2 and S3) using REVIGO (Supek et al., 2011). Figure 3 shows the enriched biological process (BP) GO-terms at different time points for cells exposed to 55 and 100 PSU, except for the 100 PSU cells at 24 h where the BP terms were not statistically enriched. Therefore, a molecular function GO-term was used to describe the acclimation path. The interaction map of enriched GO-terms clearly shows that *M. gaditana* exposed to 55 and 100 PSU use a distinct strategy to overcome salinity, except at 1 h after the transfer to the elevated salinity, where the GO-term for translation is enriched in both 55 and 100 PSU. The initial response to changes in salinity levels was significantly enriched by the photosynthesis GO-term in cells in both 55 and 100 PSU. Similarly, proteomic analysis on a halotolerant alga *Dunaliella salina* revealed the dynamics of expression of essential proteins involved in photosynthesis at high salinity conditions (Wei et al., 2017).

**Figure 3.**
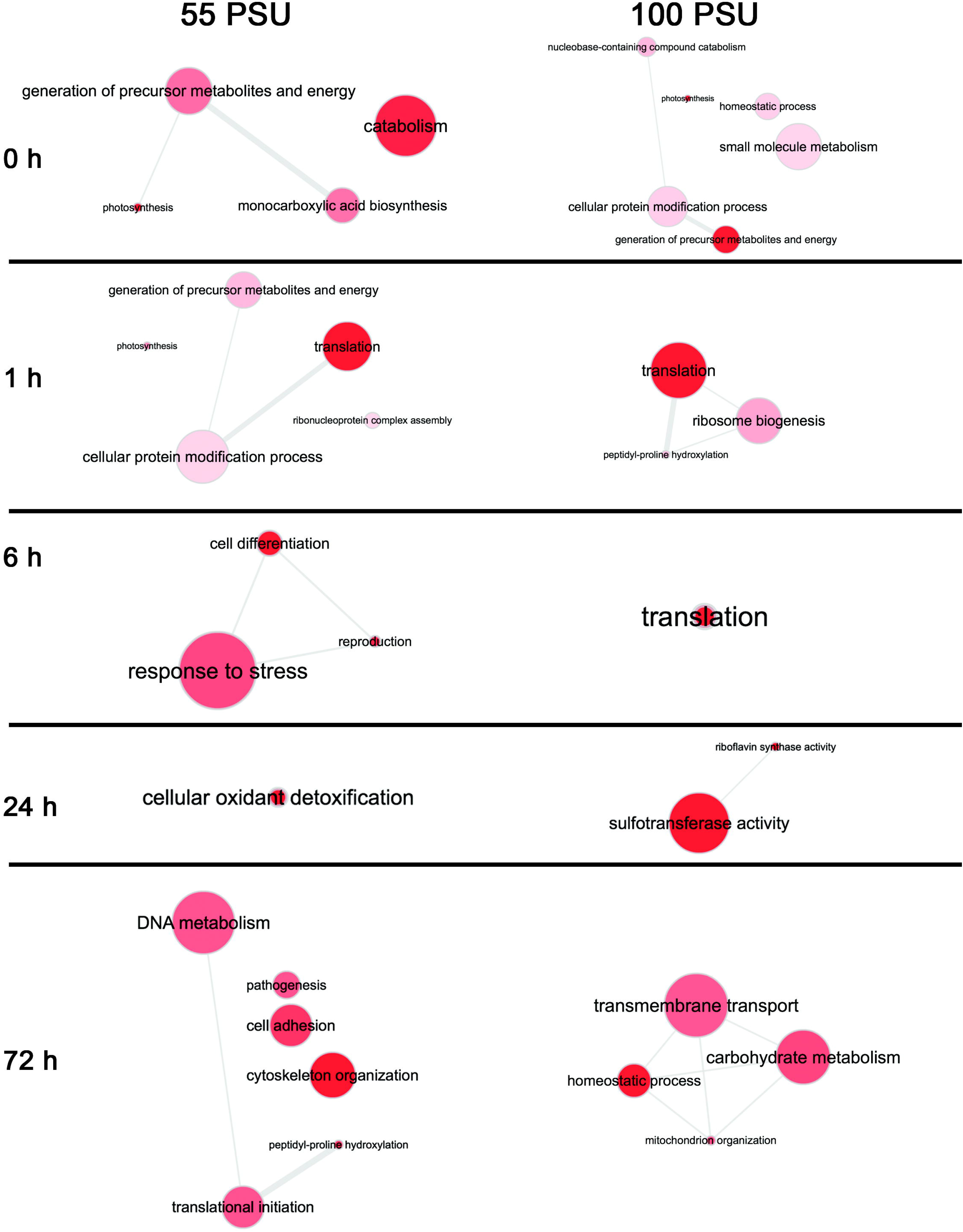
Interaction map of enriched GO-terms at different cultivation time points and salinities in *M. gaditana*. Bubble color indicates the p-value; bubble size indicates the frequency of the GO term.

Following our previous study on *M. gaditana* during the transfer to high-salinity conditions (Karthikaichamy et al., 2018), GO-terms for carbohydrate metabolism were enriched after 72 h in 100 PSU. Relatively fewer GO-terms were enriched at 6 and 24 h after transfer, which is a reflection of the number of differentially expressed proteins in cells growing at 55 and 100 PSU. However, a varied response to different saline conditions was evident across the time points. Salinity-induced ROS accumulation in cells at 55 PSU was observed in our previous study (Karthikaichamy et al., 2018) and correspondingly, the GO-terms for response to stress (GO:0033554) and cellular oxidant detoxification (GO:0098869) were enriched at 6 and 24 h respectively.

In contrast, GO-terms for energy-intensive translation (GO:0006412) was enriched at 1 and 6 h after transfer of cells to 100 PSU, suggesting a protein aided metabolic shift during the later phases (72 h) of acclimation through carbohydrate metabolism (GO:0005975) and homeostatic processes (GO:0042592). The enrichment of the carbohydrate metabolism (GO:0005975) term during the later-phase of growth suggests carbon shunting towards lipid accumulation, which is evident from our physiological (Karthikaichamy et al., 2018) and transcriptomic studies on *M. gaditana* (un-published).

### Salinity-responsive proteins (SRPs) identified by correspondence analysis

Correspondence analysis (CA) was performed to find the association between different time points and differentially expressed proteins in response to saline conditions (55 and 100 PSU). CA is a multivariate method that is used to reduce the dimensions of a complex dataset into a low-dimensional space while preserving the information (Greenacre, 1984). CA has been widely used in exploring the interactions between the genes and experimental conditions (Fellenberg et al., 2001) or tissue types (Kishino & Waddell, 2000). The temporal proteomic data was subjected to CA using ‘*ade4*’ package in R (Dray & Dufour, 2007).

The data was spread out on a two-dimensional biplot, which accounted for 73.5% and 76.5% of total inertia in cells exposed to 55 and 100 PSU respectively. In the biplot (figure 4), each point represents a protein, and the arrows that originate from origin represent time points. Proteins closer or along the arrow are up-regulated at that particular time point, and the proteins that are on the opposite quadrant of the arrow represent the down-regulated protein at that particular time point. Proteins are colored according to the contribution towards the plot; the highest contribution indicates that the protein has been highly expressed at that particular time point.

**Figure 4.**
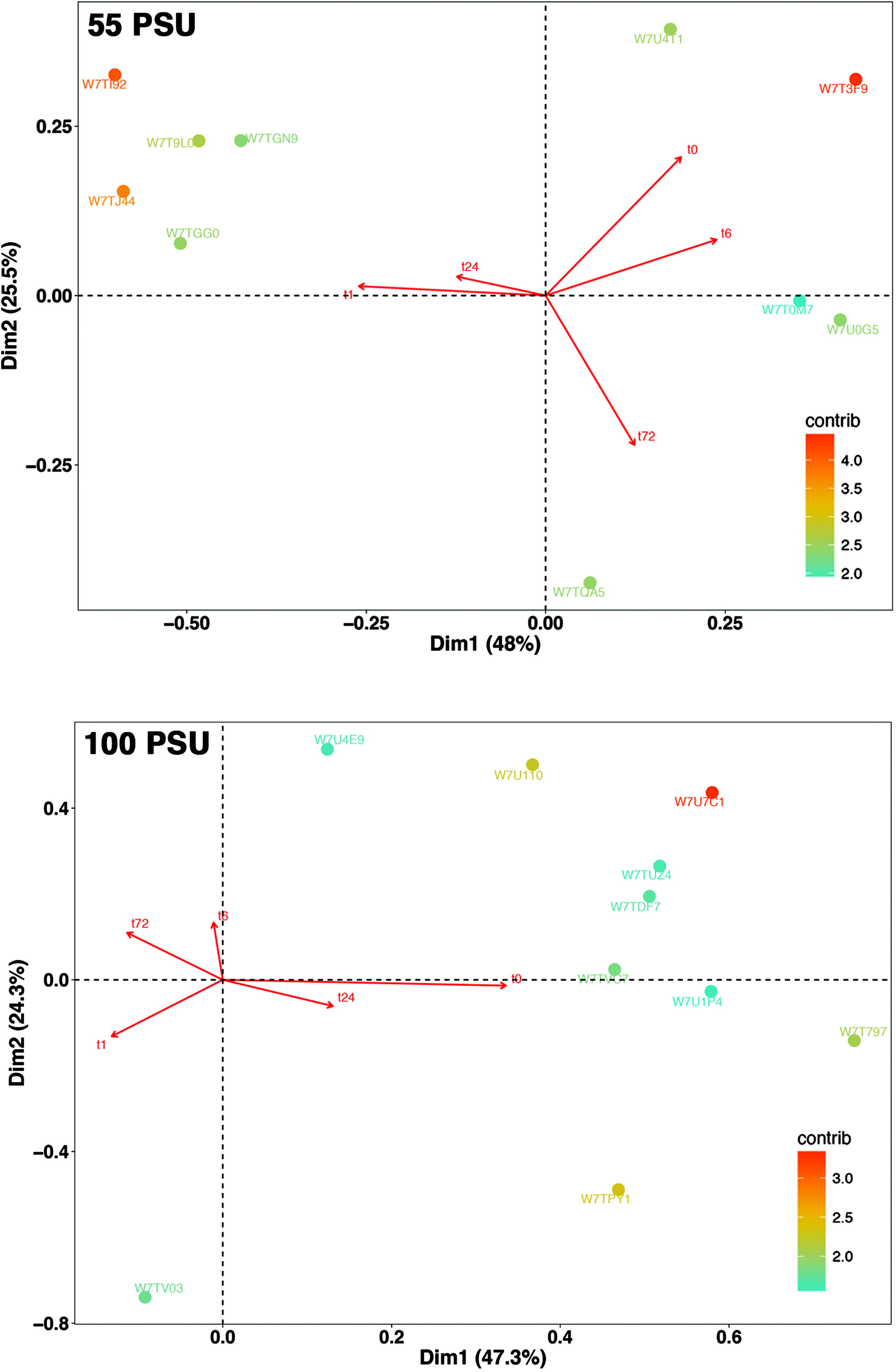
Correspondence analysis of differentially expressed proteins in *M. gaditana* at different time points in 55 and 100 PSU. Biplot showing the association between time points and differentially expressed proteins (numbers along the red arrow represent time points).

### Top salinity-responsive proteins in cells grown in hypersaline conditions

Top 10 proteins, based on their contribution values, for each salinity conditions, are shown in figure 5. Three uncharacterized proteins were found to be among the top 10 contributors in 55 PSU (W7T9L0, W7TQA5, and W7T0M7) and two were found in cells exposed to 100 PSU (W7U7C1 and W7T797). The top contributor in cells exposed to 55 PSU was W7T3F9, an esterase lipase thioesterase family protein, which was up-regulated at 0 h and 6 h. It has been GO annotated with acylyglycerol lipase activity (GO:0047372) and medium-chain fatty acid catabolic processes (GO:0051793). The gene product (at2g39420) of the protein homolog in *Arabidopsis* (Q8RXN7_ARATH; E-value: 1.7e-65) was up-regulated (1.7 log2FC) in response to salt and heat stress. It also responds to drought stress and ozone exposure (Abdeen et al., 2010; Suzuki et al., 2016). The esterase lipase thioesterase family protein is widely involved in mobilizing oil reserves to provide carbon backbones for energy sources during abiotic and heavy metal stress (Gao et al., 2010; Troncoso-Ponce et al., 2013). The other early response protein is Nudix hydrolase 6-like protein (W7U4T1), which was up-regulated ~64-fold at 0 h. The *Arabidopsis* Nudix hydrolases positively regulate salicylic acid-mediated signaling and modulates defense responses against abiotic and biotic stress (Ge & Xia, 2008; Jambunathan & Mahalingam, 2006).

**Figure 5.**
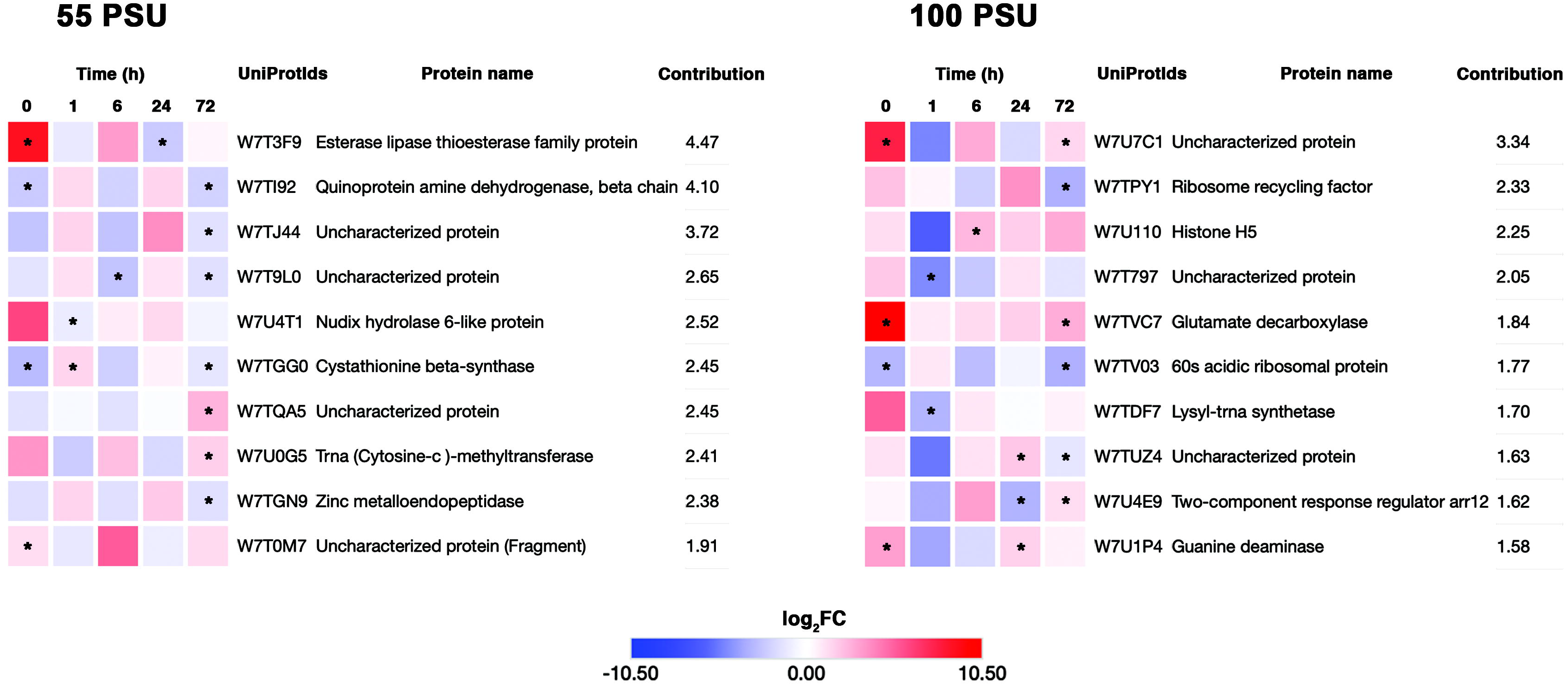
Heatmap of the top 10 differentially expressed proteins in *M. gaditana* based on the contribution towards the dimensions in the biplot. Color scale indicate log_2_FC value. Significant levels of protein abundance are marked with an asterisk.

Another protein whose abundance is highly modulated during the high saline condition (55 PSU) is quinoprotein amine dehydrogenase, beta chain (W7TI92). W7TI92 was up-regulated ~3-fold at 1 and 24 h, and down-regulated up to 5.5-fold at other time points (0, 6 and 72 h). The quinoprotein amine dehydrogenase, beta chain has a beta-propeller structure, which is similar to the YVTN (Tyr-Val-Thr-Asn) repeat in the N-terminal of archaeal surface proteins. The structural similarity to the archaeal surface proteins is of high significance because it confers protection to the cell during extreme environmental conditions (Jing et al., 2002). Cystathionine beta-synthase (W7TGG0) which catalyzes the formation of cystathione from L-serine and L-homocysteine is variably expressed through-out the growth phase and the highest expression observed at 1 h (~3x). The ortholog of W7TGG0 in rice plant (OsCBSCBSPB4), when over-expressed, was found to be responsible for the increase in biomass and delayed leaf senescence during extreme conditions (high saline, dehydration, high temperature) (Kumar et al., 2018).

An ortholog of W7T0M7 (Uncharacterized protein (Fragment)) was up-regulated ~32-fold at 6 h, and a modest down-regulation and up-regulation were observed at 1, 24 h and 0, 72 h respectively. The ortholog (A0A0A7CLB7) was found in a fungus *Thraustotheca clavate* (E-value: 8.4e-9) is a secreted protein, belonging to RNase T2 family (IPR033697). Plant T2 RNases scavenge phosphate from ribonucleotides during senescence, and they also respond to wounding or pathogen invasion (Deshpande & Shankar, 2002).

On the other hand, the top contributing protein in 100 PSU is an uncharacterized protein (W7U7C1), which is predicted to be localized in chloroplast, mitochondria, and nucleus. The protein is found in high abundance (~215x) during the initial phases (0 h) of growth in 100 PSU and at a modest up-regulation of up to 8x and 3X at 6 and 72 h respectively. The paralog for W7U7C1 is an Hsp70-interacting protein (K8YSA6; E-value: 7.8e-133), which is involved in Hsp70 stabilization in ADP (adenosine diphosphate) bound conformation (Palleros et al., 1993). Hsp70-interacting protein (Hip), which is ubiquitously found in animals and plants is involved in the regulation of Hsp70 assisted protein folding and import (Boorstein et al., 1994; Boston et al., 1996; Miernyk, 1999). The strong up-regulation of W7U7C1 during the initial stages of growth in 100 PSU signifies the importance of protein folding upon stress-induced protein unfolding, thereby maintaining proteostasis during stress conditions (Kim et al., 2013). One of the most abundant proteins expressed in response to the high saline condition is glutamate decarboxylase (W7TVC7). It was strongly up-regulated by 1450-fold at 0 h, and then a modest up-regulation (up to 8x) was observed at other time points. Plants produce Y-aminobutyrate by irreversible α-decarboxylation of glutamate by glutamate decarboxylase. GABA acts as a natural defense against pests, and it is also associated with stress response in tobacco and *Camellia sinensis* (tea) plants (McLean et al., 2003; Mei et al., 2016).

Proteins involved in translation were variably affected at the highest saline conditions (100 PSU). For instance, ribosome recycling factor (W7TPY1) and 60s acidic ribosomal protein (W7TV03) responded well to change in salinity levels. While the ribosome recycling factor protein (W7TPY1), which is involved in translation and chloroplast biogenesis was up-regulated up to 12-fold at 0 and 24 h, the 60 S acidic ribosomal protein (W7TV03), which plays an essential part in elongation of proteins (Naganuma et al., 2010) was consistently down-regulated (up to ~11x) at all time points, except at 1 h where a modest up-regulation (~1.8) was observed. The availability of nitrogen could limit the translation of protein, and therefore, plants have evolved a strategy to salvage nitrogen from an N rich source such as ureides. Guanine deaminase (W7U1P4) is an enzyme involved in purine nucleotide catabolism, in which guanine is converted into xanthine and then to allantoin in further steps of catabolism, which is considered to be the first step of salvaging nitrogen from guanine. Allantoin accumulation is linked with salt stress tolerance in *Arabidopsis thaliana* (Irani & Todd, 2016; Lescano et al., 2016). Likewise, in our study, the levels of guanine deaminase (W7U1P4) was up-regulated (up to ~8x) at 0, 24 and 72 h, which indicates a possible salvage mechanism of nitrogen (N2) by *M. gaditana* from purine nucleotides.

### Salinity induced proteome changes in cellular metabolism

The proteome dynamics of *M. gaditana* was studied under salinity stress conditions. The differentially expressed proteins at various time points were subjected to GO enrichment analysis, which revealed a diverse response to various high salinity levels (55 and 100 PSU). Similarly, we observed salinity induced dynamics in the transcriptome of *M. gaditana* (un-published). In order to understand the fate of cellular metabolism at high saline conditions, and to connect a line between physiology (Karthikaichamy et al., 2018) and transcriptome (un-published) of salinity induced *M. gaditana*, the differentially expressed proteins were analyzed for their biological significance in the following sections.

### Mitigation of ROS levels by rapid accumulation of stress response proteins

Imbalance in light-harvesting and utilization in salinity stressed phototrophs could generate ROS, which is deleterious to the cell (Miller et al., 2010). ROS scavenging and homeostasis are critical for the shift in metabolism after salinity stress is imposed. Several proteins involved in oxidative stress responses were up-regulated at the high saline conditions (55 and 100 PSU) (supplementary table S4). Overall, the fold change of proteins involved in oxidative stress responses (such as superoxide dismutase, catalase, and ferredoxin) was up-regulated (up to ~8x) at 1 h in 55 PSU and 1 h and 72 h in 100 PSU, except a few proteins such as superoxide dismutase (W7TM73) and ferredoxins (W7TM71). The other superoxide dismutases (W7T7U7, W7TGS6, W7U428) were significantly expressed (up-regulation up to ~5x at 1 and 72 h) in 100 PSU. Several glutaredoxins, glutathione transferase (W7TNY9; W7U7N8), ascorbate peroxidase (W7U0U0) and thioredoxins were significantly altered in 100 PSU. ROS accumulation is a signature of stress response in plants and microalgae. Comparable with the metabolic changes, the salinity induced ROS accumulation triggered a set of ROS scavenging proteins such as SOD, CAT, glutathione peroxidase, ferredoxin in *M. gaditana*. These proteins are involved in maintaining the redox status in higher plants (Chen et al., 2009; Kav et al., 2004; Sugimoto & Takeda, 2009). Likewise, the SOD level was increased in the halotolerant alga *Dunaliella salina* (Wei et al., 2017) and in a salinity tolerant strain of *Chlamydomonas reinhardtii* (Sithtisarn et al., 2017). Anti-oxidant proteins help to maintain the ROS level inside the cell, and accumulation of such proteins have been found in salinity treated plants (Miller et al., 2010). Mitigation of ROS levels by rapid accumulation of scavenging enzymes could be an initial step towards acclimatization of *M. gaditana* to saline conditions. The expression profiles of the significantly altered oxidative stress response proteins are shown in Figure 6.

**Figure 6.**
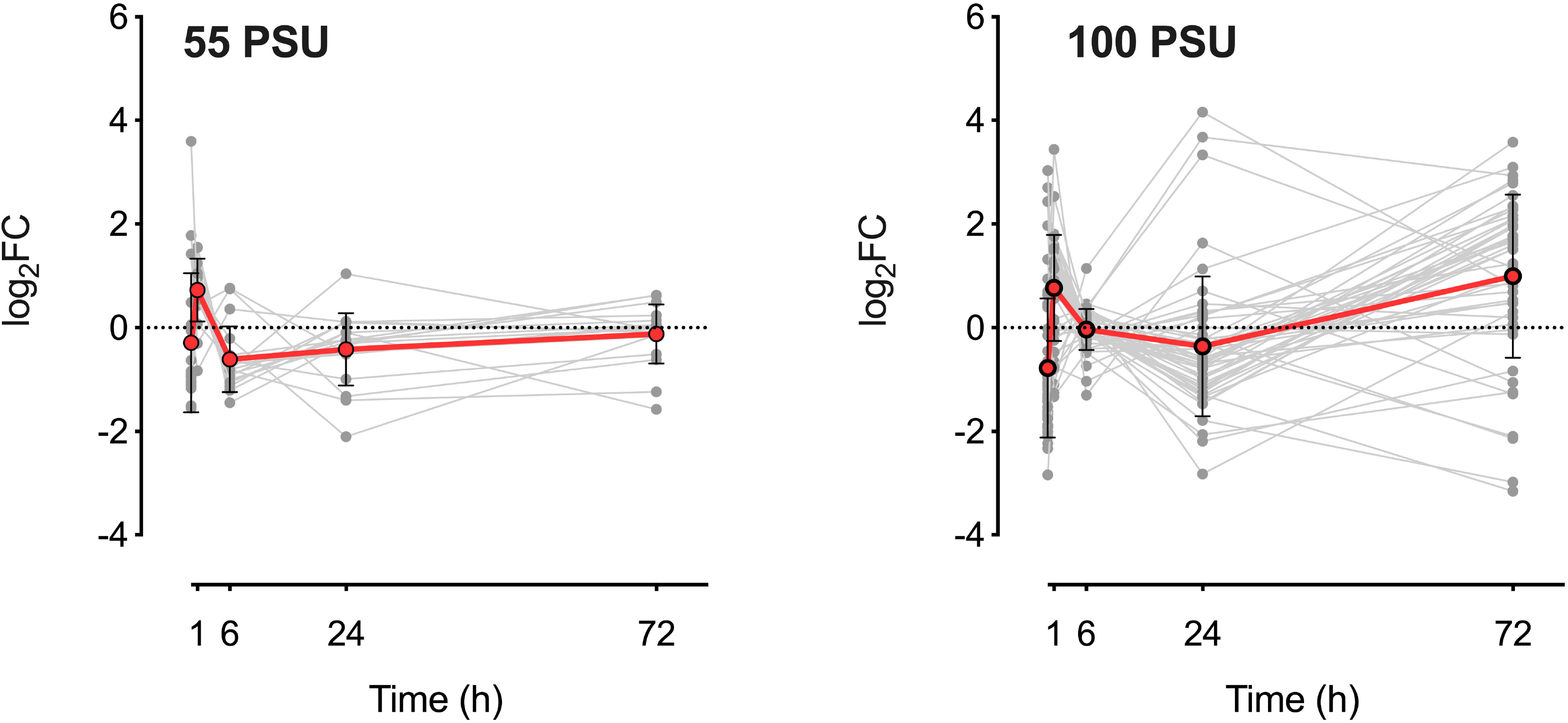
Expression levels of proteins involved in oxidative stress responses in *M. gaditana* (grey lines). The red line indicates the mean expression values of the significantly altered proteins. Error bars indicate standard deviations.

### Hypersalinity induces up-regulation of osmolyte biosynthesis proteins

Osmolyte production is one of the primary responses of microalgae towards an increase in salinity (Flowers & Colmer, 2008). The osmolytes are highly soluble, low-molecular-weight molecules primarily accumulated in the cytoplasm. They range from simple carbohydrates such as trehalose, xylose and sucrose to amino-acids such as glycine, proline, or amino-acid derivatives such as ornithine (Flowers et al., 1977; Wyn Jones, 1977). The expression profiles of the proteins involved in osmolyte biosynthesis are shown in Figure 7. In our study, we observed an increase in protein expression levels of key enzymes involved in trehalose, glycine betaine, and glycerol biosynthetic pathways (supplementary table S5).

**Figure 7.**
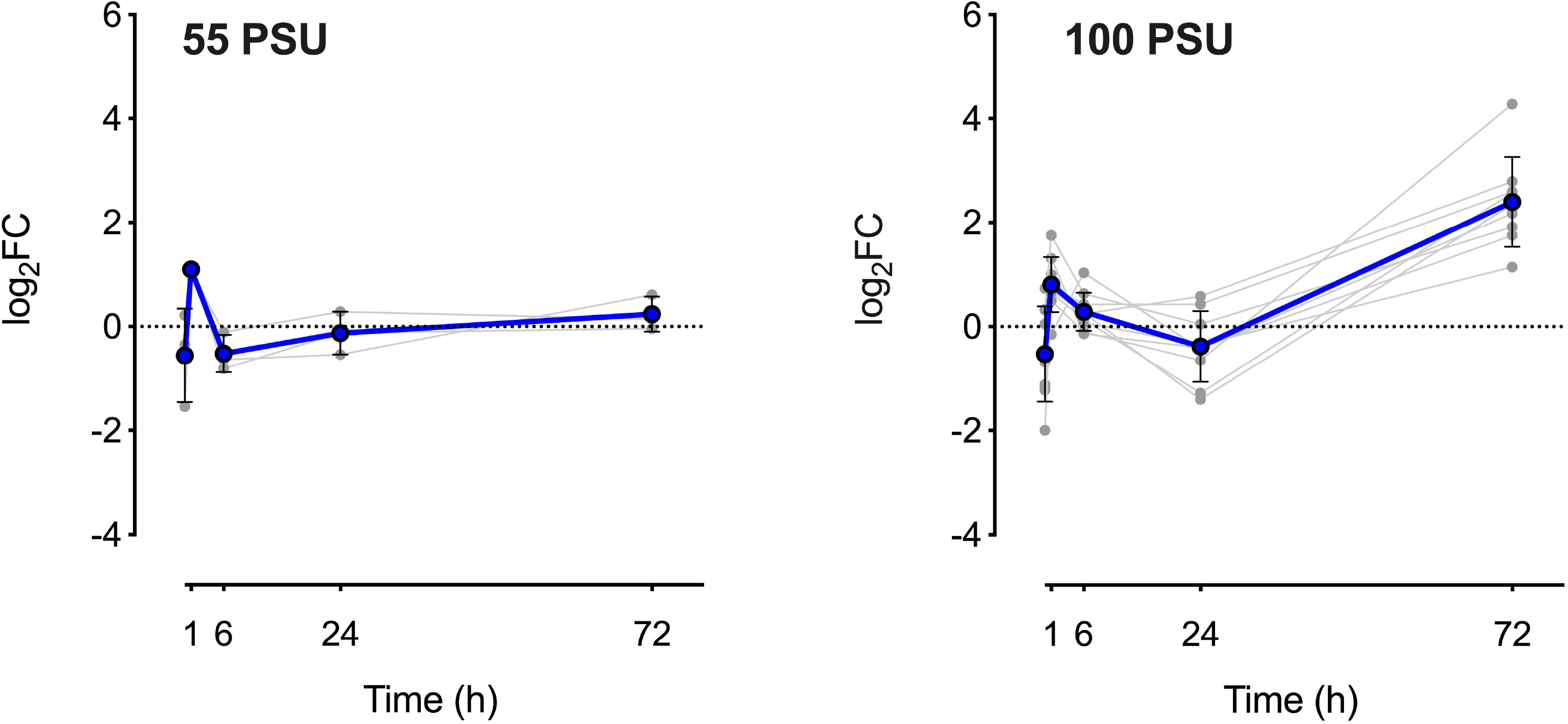
Expression levels of key enzymes involved in osmolyte biosynthetic pathway in *M. gaditana* under high saline conditions (grey lines). Blue line indicates the mean expression values of the significantly altered proteins. Error bars indicate standard deviations.

Trehalose is a non-reducing disaccharide formed by two glucose molecules linked by alpha-1,1-bond. Accumulation of trehalose during stress conditions such as drought, salt, heat, desiccation and freeze stress has been reported across various microorganisms, including plants and microalgae (Goddijn & van Dun, 1999; Hershkovitz et al., 1991; Hirth et al., 2017). Moreover, trehalose confers tolerance against abiotic stresses in rice (Garg et al., 2002) and it is found to be essential for carbohydrate utilization in *Arabidopsis thaliana* (Schluepmann et al., 2003). The levels of trehalose-6-phosphate synthase (W7TQF4), trehalose phosphate synthase (W7TD30 and W7TBU1) and trehalose 6-phosphate phosphatase (W7TFM0) were significantly altered only in 100 PSU, and they were up-regulated (up to ~16x) during the later phases of growth (72 h).

The enzyme that prime trehalose biosynthetic pathway, UDP-glucose pyrophosphorylase 2 (W7TFM0), by providing UDP-glucose as substrate was significantly altered only in 100 PSU. The protein abundance increased (~19x) at 72 h in100 PSU. Interestingly, the expression pattern of UDP-glucose pyrophosphorylase 2 is similar to the trehalose biosynthetic proteins, implying the importance of the UDP-glucose pool for trehalose biosynthesis during high salinity conditions (Figure 8).

**Figure 8.**
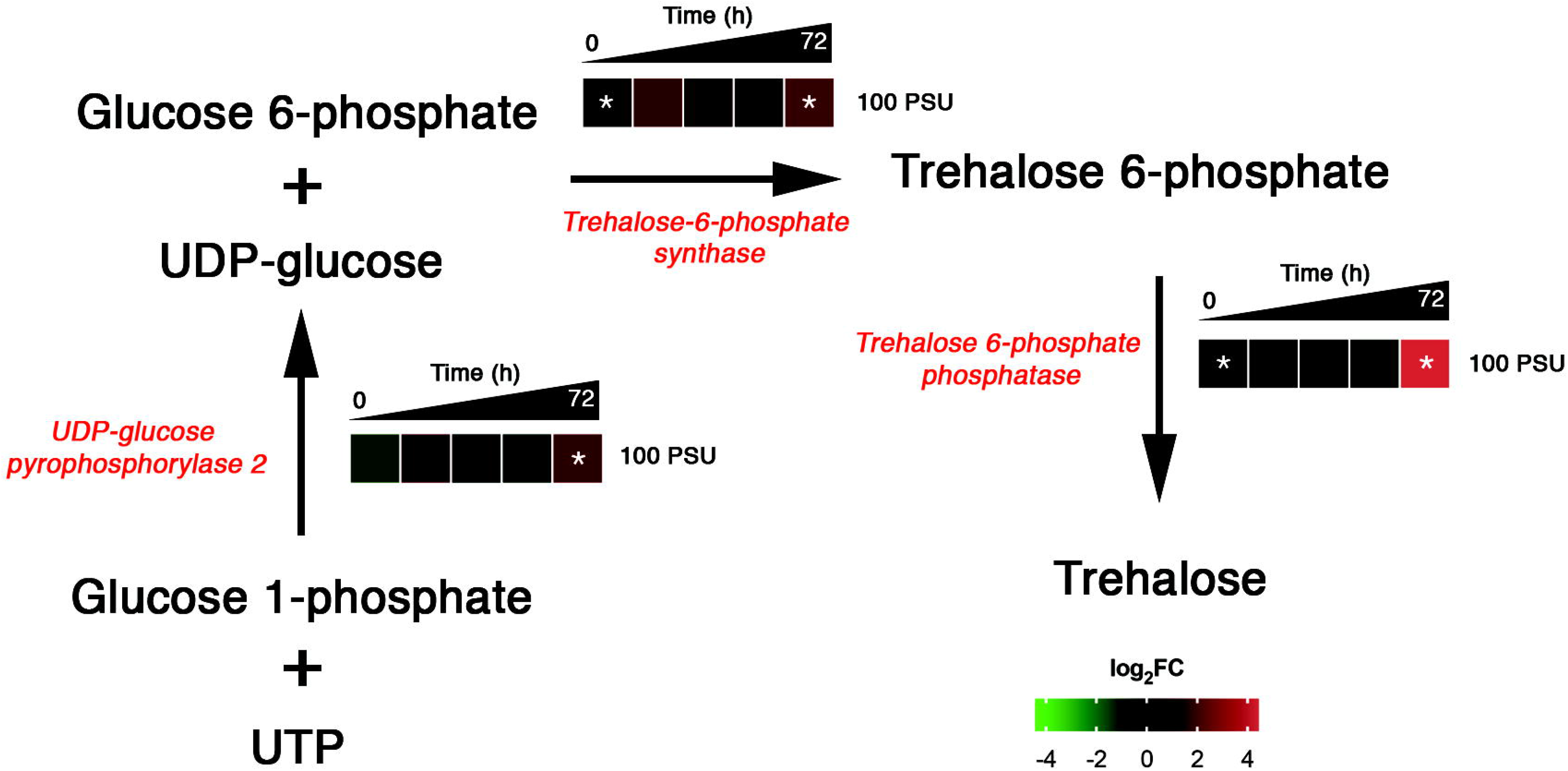
Salinity induced temporal dynamics of proteins involved in trehalose biosynthesis in *M. gaditana*. Significant levels of protein abundance are marked with an asterisk.

Other proteins involved in compatible solute biosynthesis include choline dehydrogenase (W7TDH7, W7TN63), which catalyzes the first step in glycine betaine biosynthesis through betaine aldehyde, and which was significantly altered only in 100 PSU. The maximum up-regulation (up to ~4x) occurred at 1 and 72 h post salinity stress induction. Similar dynamics in betaine biosynthesis was observed in our earlier transcriptomic study. The assimilation of CO_2_ into glycine betaine during saline conditions was observed in *A*. *halophytica* (Takabe et al., 1988). Also, exogenous addition of glycine betaine to the growth medium of *A. thaliana* conferred tolerance to abiotic stresses (Hibino et al., 2002).

A key enzyme in glycerol (compatible solute) biosynthesis, glycerol-3-phosphate dehydrogenase (W7TIR6) is significantly up-regulated (up to ~8x) during later phases of growth (72 h) in 100 PSU. GPDH is located in the mitochondrial inner-membrane space or cytosol and catalyzes the reduction of dihydroxyacetone phosphate into glycerol-3-phosphate. It also links carbohydrate and lipid metabolism by providing glycerol backbones for triacylglycerol (TAG) synthesis (Ou et al., 2006). Salinity induced *Dunaliella salina* accumulates a significant amount of glycerol, which is comparable with GPDH activity (Wei et al., 2017).

### Proteins involved in fatty acid metabolism

Fatty acids are energy reservoirs of the cell that can be used during nutrient limitation conditions or any adverse environmental conditions (Mata-Pérez et al., 2016). Cell membranes play a vital role in sensing the stress conditions and initiate the acclimatization reactions. The fluidity of plant and algal cell membrane can be altered by changing the fatty acid composition of the membrane. This phenomenon is beneficial for the organism to adapt to the stress conditions (Mikami & Murata, 2003; Wada et al., 1994). Salt induced changes in neutral-lipid content in *M. gaditana* have already been reported in our previous study (Karthikaichamy et al., 2018). Comparable levels of transcript induction of lipid biosynthetic genes were observed in *M. gaditana*. The expression profiles of significantly altered proteins during hypersaline conditions are shown in Figure 9.

**Figure 9.**
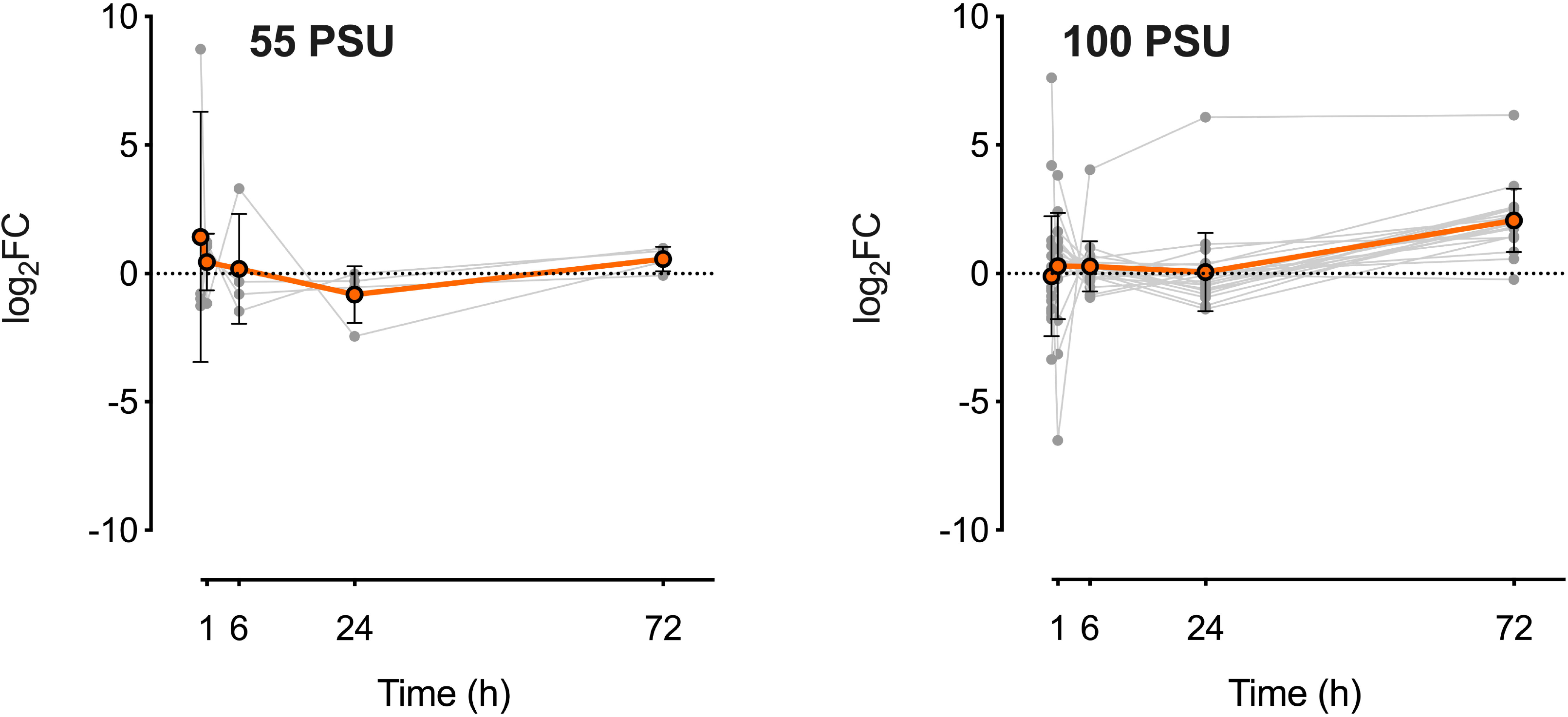
Expression levels of proteins involved in fatty acid metabolism (grey lines) in *M. gaditana*. Orange line indicates the mean expression values of the significantly altered proteins. Error bars indicate standard deviations.

In line with our physiological analysis, we have identified proteins involved in lipid and fatty acid metabolism (supplementary table S6). Four proteins involved in lipid metabolism were statistically altered in 55 PSU, among which are an esterase lipase thioesterase family protein (W7T3F9), fatty acid desaturase type 2 (W7UCJ8) and 3-oxoacyl-(Acyl-carrier-protein) reductase (W7U8F0), which were up-regulated more than 2-fold. 3-oxoacyl-(Acyl-carrier-protein) reductase (W7U8F0) is involved in the NADPH-dependent reduction of beta-ketoacyl-ACP substrates in the elongation cycle of fatty acid biosynthesis (Lai & Cronan, 2004). An instantaneous response to salinity stress was observed with an abundance of esterase lipase thioesterase family protein (W7T3F9) during the early phases of growth (0 h) in both saline conditions (55 and 100 PSU). The implications of esterase lipase thioesterase family protein during the alleviation of salinity stress have been discussed earlier.

In 100 PSU, proteins involved in lipid biosynthesis such as long-chain acyl-CoA synthetase (W7TGG5), diacylglycerol acyltransferase family protein (W7U9S5), monogalactosyldiacylglycerol synthase, long-chain acyl-synthetase 7 (W7TAS6) and several lipases were statistically up-regulated (up to ~8x) during the later phases of growth (72 h). The significant up-regulation of the proteins involved in lipid metabolism is consistent with our previous physiological studies on *M. gaditana* (Karthikaichamy et al., 2018). Long-chain acyl-CoA synthetase (LACS) is indispensable for biosynthesis of fatty acid derived molecules. In germinating *Arabidopsis thaliana* seedlings, LACS activate the free fatty acids to the β-oxidation pathway for the removal of two-carbon molecule in the form of acetyl-CoA (Fulda et al., 2004). The acetyl-CoA thus formed enters the TCA cycle for the production of cellular energy (Gerhardt, 1992). It is therefore evident that *M. gaditana* prefers β-oxidation of accumulated fatty acid for sustaining the energy requirements during high saline conditions. Though the cell primarily focusses on β-oxidation of lipids, a comparable up-regulation (~8x) of diacylglycerol acyltransferase family protein (W7U9S5), which is involved in triacylglycerol (TAG) synthesis is observed during the later phases (72 h) in 100 PSU. This phenomenon could account for a steady, but a not sharp, increase in neutral lipid content in *M. gaditana* (Karthikaichamy et al., 2018). *Arabidopsis thaliana* diacylglycerol acyltransferase mutants were sensitive to several stress conditions, implying a possible role in stress mitigation (Lu & Hills, 2002). Lipolytic protein g-d-s-l family protein (W7TVB2) was significantly up-regulated (up to ~64x) at later-phases of growth in 100 PSU. The lipolytic protein g-d-s-l family have been studied in plant species such as *Arabidopsis*, rice, and maize, and it has been associated with plant development and defense responses (Brick et al., 1995). Overexpression of GDSL◻motif lipase increases tolerance in transgenic *A. thaliana* (Naranjo et al., 2006).

### Oxidative pentose-phosphate pathway redirects the carbon flux towards fatty acid biosynthesis under high saline conditions

The primary source of NADPH for desaturation and elongation reactions of polyunsaturated fatty acid (PUFA) is malic enzyme (ME). However, in our study, malic enzyme was not significantly up-regulated. Various studies have established a connection between malic enzyme levels and neutral lipid accumulation (Zhu et al., 2018). Also, the transcript abundance of malic enzyme was down-regulated in our transcriptomic analysis on *M. gaditana* (un-published). Corresponding to the previous hypothesis established on the use of oxidative pentose phosphate pathway as an alternative source to TCA cycle for reducing equivalents, one of the key proteins (glucose-6-phosphate 1-dehydrogenase) involved in the pentose phosphate pathway was significantly up-regulated (~16x) during the later phases of growth (24 and 72 h) in 100 PSU (supplementary table S6). Glucose-6-phosphate 1-dehydrogenase (W7TZ08), catalyzes the rate-limiting step of the oxidative pentose-phosphate pathway (PPP) and provides reducing equivalents (NADPH) and pentose phosphates for fatty acid and nucleic acid biosynthesis respectively (Wang et al., 2014). Thus, *M. gaditana* establishes lipid homeostasis via simultaneous activation of β-oxidation and TAG biosynthesis proteins during high saline conditions. Another rate-limiting enzyme in PPP, 6-phosphogluconate dehydrogenase (PGDH), decarboxylating (W7TN92) was significantly up-regulated (up to ~16x) during the later phases of growth (72 h), suggesting that PPP is enhanced during high saline condition in *M. gaditana.* PPP is a key pathway in carbohydrate metabolism in plants and links other biosynthetic processes such as lipid, nucleotide, sugar and aromatic amino acid biosynthesis by providing reducing equivalents (NADPH) and precursor molecules. The increase in abundance of the critical proteins involved in reductive pentose phosphate pathway complements the increased glycerol biosynthetic protein abundance. PPP aids in glycerol accumulation by replenishing the carbon intermediates (ribulose-5-P). A similar trend was observed in higher plants (rice and wheat), where salt stress induced the expression of PGDH and GPDH (Krishnaraj & Thorpe, 1996). Our results suggest that PPP helps in redirecting the carbon flux towards fatty acid biosynthesis under high saline conditions in *M. gaditana*.

### Photosynthesis and redox energy metabolism

Salinity stress is often linked with the reduction in photosynthetic efficiency (Karthikaichamy et al., 2018), primarily due to damage caused by ROS to light-harvesting complexes (LHCs) and oxygen evolution factor proteins of PSI (Subramanyam et al., 2010). In other instances, cessation of PSII activity was observed in *Dunaliella tertiolecta* and *Nannochloropsis* sp. under high salinity conditions (Gilmour et al., 1985; Martínez-Roldán et al., 2014). A comprehensive list of differentially expressed proteins involved in photosynthesis is provided in supplementary table S7.

Porphobilinogen deaminase (W7UC09), which catalyzes the primary step of chlorophyll biosynthesis (Roberts et al., 2013) was down-regulated (up to ~4x) throughout the growth phase (significant down-regulation at 72 h), except at 1 h. The reduction in the abundance of a critical chlorophyll biosynthetic protein was reflected in the reduction of chlorophyll content in salinity stressed *M. gaditana* (Karthikaichamy et al., 2018). Reduction in chlorophyll biosynthetic protein levels was also observed in salinity stressed *D. salina* (Wei et al., 2017). Reduction in chlorophyll levels during stress conditions can be associated either with the cessation of biosynthesis or dilution by cellular growth. The latter was reported in *C. reinhardtii* (Juergens et al., 2015), while the former phenomenon was observed in our case. Interestingly, various proteins associated with PSII (oxygen evolving complex, D1 protein, D3 protein, PsbP, cytochrome *c*, ferredoxin-NADP reductase, PSII 47 kDa protein and light-harvesting complex proteins) and PSI (apoprotein A1, A2 and PSI reaction center proteins) were up-regulated (up to ~8x) during the initial phases of growth (0 and 1 h) in both the salinity levels (55 and 100 PSU). A few PSII and PSI proteins were also up-regulated (up to ~8x) during the later phases of growth in 100 PSU. The inner antenna PSII 47 kDa protein is conserved and is essential for assembling PSII in algae and higher plants (Summer et al., 1997). Cytochrome *b*559 subunit alpha (K9ZWI6) and cytochrome *b*559 subunit beta (K9ZVG9) protects PSII from photo-inhibition by enhancing cyclic electron transport (CET) in *Chlamydomonas reinhardtii* (Burda et al., 2003) is significantly up-regulated (up to ~2x) in 55 PSU. Enhanced photosynthesis and energy metabolism during high salinity conditions has been linked with increased CO_2_ fixation and glycerol biosynthesis in *Dunaliella salina* (Liska et al., 2004). The expression profiles of the proteins involved in photosynthesis are shown in Figure 10.

**Figure 10.**
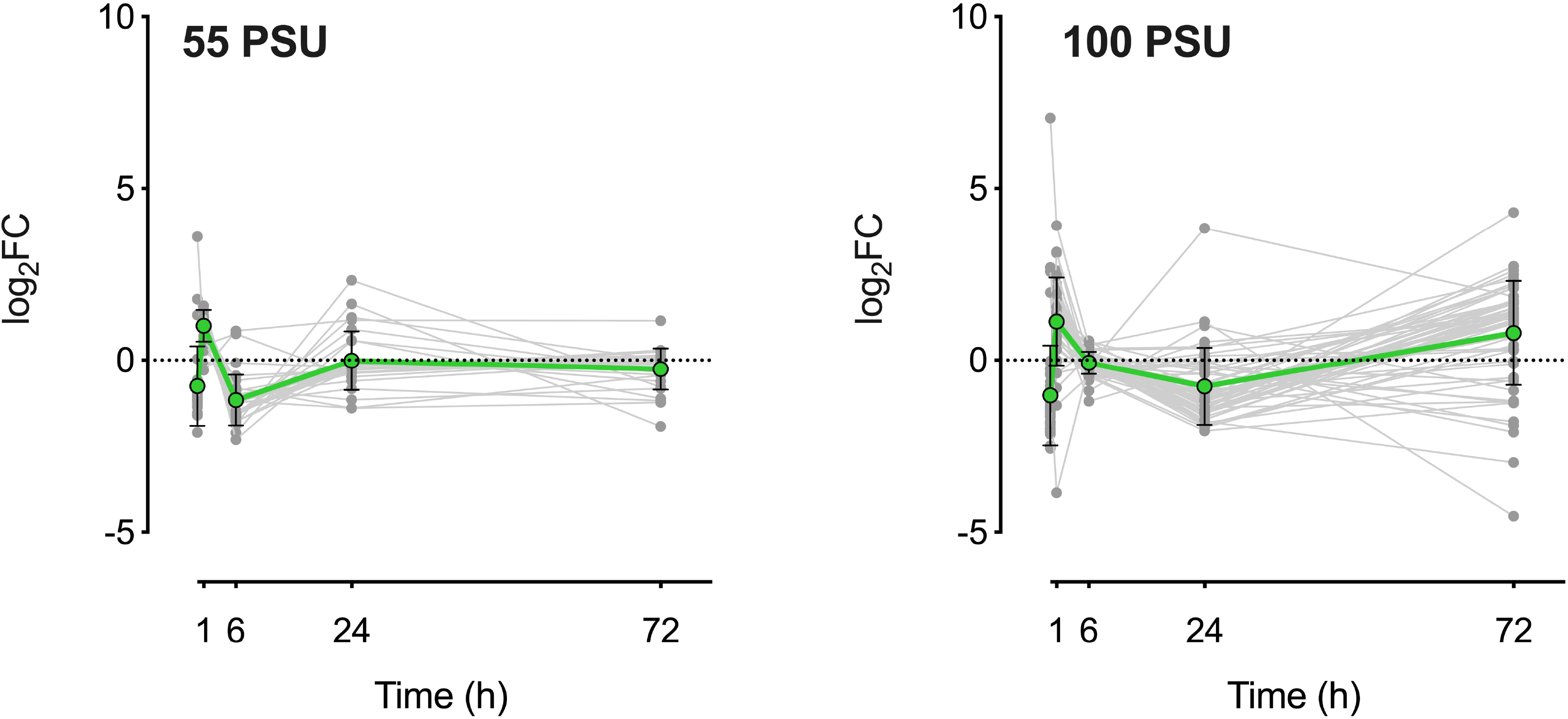
Expression levels of proteins involved in photosynthesis (grey lines) in *M. gaditana*. Green line indicates the mean expression values of the significantly altered proteins. Error bars indicate standard deviations.

Several light-harvesting complexes of PSII (LHCII) were found to be up-regulated (up to ~5x) in 55 (1 h) and 100 PSU (1 and 72 h). Enhancement of LHCII proteins was also observed upon palmella formation in salinity induced *Dunaliella salina*. During stress conditions, regulation of redox homeostasis in the chloroplast is enhanced by reversible phosphorylation of LHCII proteins (Grieco et al., 2012). In photosynthetic micro-organisms and higher plants, carbonic anhydrases (CA) are responsible for converting HCO_3_^−^ to CO_2_, which is then fixed by ribulose 1,5-bisphosphate carboxylase/oxygenase (Espie & Kimber, 2011). There are multiple forms of carbonic anhydrases involved in a host of reactions in the cell, and an external carbonic anhydrase that plays a role in CO_2_ concentrating mechanisms. Up-regulation (up to ~8x) of carbonic anhydrase (W7TZZ4) was observed only in 100 PSU at 72 h. The protein abundance of the target of carbon concentrating mechanism (CCM), ribulose bisphosphate carboxylase (K9ZV74, K9ZWI1) was not significant. Thus, the protein abundance of carbonic anhydrase could be beneficial for the cell during adverse conditions rather than to invest in an energetic input for Rubisco biosynthesis (Young & Hopkinson, 2017). Upregulation of external CA is induced by hypersaline conditions in *Dunaliaella salina* as dissolved CO_2_ levels are lower with increasing salinity. While, Rubisco activity was inhibited in response to high saline conditions in *D. salina* (Booth and Beardall, 1991).

## Conclusions

The current study aimed to understand the temporal proteome dynamics during acclimatization to high saline conditions (55 and 100 PSU). The timepoints and salinity conditions were chosen based on our previous physiological studies on *M. gaditana* at high saline conditions. Significantly altered proteins at each time point were enriched with GO-terms using fisher exact analysis. GO-enrichment revealed the acclimatization path of *M. gaditana* towards hypersaline conditions. Salinity level of 100 PSU had significant impact on the proteome of *M. gaditana*. Correspondence analysis revealed the salinity-responsive proteins and their possible metabolic implications was analyzed. Proteins involved in lipid and osmolyte biosynthesis was observed to be up-regulated at high salinity conditions, which could be one of the possible strategies of *M. gaditana* in acclimatizing to high salinity conditions. Osmolyte helps in retaining the cellular morphology by acting as regulators of turgor pressure. Lipid biosynthetic proteins were upregulated at high salt conditions, so as the proteins involved in fatty acid oxidation. The oxidized product could serve as a precursor for various biosynthetic pathways. Protein levels and the physiology of *M. gaditana* at high salt conditions correspond to each other. This study provides a comprehensive overview of the temporal proteome level changes in *M. gaditana* at high salt conditions. A probable acclimatization strategy of *M. gaditana* during high-saline conditions was discussed, emphasizing on osmolyte, lipid and photosynthesis and redox energy metabolism pathways. The proteome analysis suggests a possible route towards acclimatization and a few potential target proteins for strain improvement.

## Acknowledgements

The authors acknowledge the technical support from the Monash Proteomics & Metabolomics Facility, Monash University, Australia. This work was supported by the IITB-Monash Research Academy, IITB, Mumbai, India; Reliance Industries Limited, India (IMURA0304).

## References

Abdeen, A., Schnell, J., Miki, B. 2010. Transcriptome analysis reveals absence of unintended effects in drought-tolerant transgenic plants overexpressing the transcription factor ABF3. in: BMC genomics, Vol. 11, pp. 69.

Aghaei, K., Ehsanpour, A.A., Komatsu, S. 2008. Proteome analysis of potato under salt stress. J Proteome Res, 7(11), 4858–68.

Boorstein, W.R., Ziegelhoffer, T., Craig, E.A. 1994. Molecular evolution of the HSP70 multigene family. J Mol Evol, 38(1), 1–17.

Booth, W., Beardall, J. 1991. Effects of salinity on inorganic carbon utilization and carbonic anhydrase activity in the halotolerant alga *Dunaliella salina* (Chlorophyta). Phycologia, 30(2), 220–225.

Boston, R.S., Viitanen, P.V., Vierling, E. 1996. Molecular chaperones and protein folding in plants. Plant Mol Biol, 32(1-2), 191–222.

Brennan, L., Owende, P. 2010. Biofuels from microalgae—a review of technologies for sproduction, processing, and extractions of biofuels and co-products. Renewable and sustainable energy reviews, 14(2), 557–577.

Brick, D.J., Brumlik, M.J., Buckley, J.T., Cao, J.X., Davies, P.C., Misra, S., Tranbarger, T.J., Upton, C. 1995. A new family of lipolytic plant enzymes with members in rice, *Arabidopsis* and maize. FEBS Lett, 377(3), 475–80.

Burda, K., Kruk, J., Borgstadt, R., Stanek, J., Strzalka, K., Schmid, G.H., Kruse, O. 2003. Mossbauer studies of the non-heme iron and cytochrome b559 in a *Chlamydomonas reinhardtii* PSI-mutant and their interactions with alpha-tocopherol quinone. FEBS Lett, 535(1-3), 159–65.

Chen, S., Gollop, N., Heuer, B. 2009. Proteomic analysis of salt-stressed tomato (*Solanum lycopersicum*) seedlings: effect of genotype and exogenous application of glycinebetaine. Journal of Experimental Botany, 60(7), 2005–2019.

Conesa, A., Gotz, S., Garcia-Gomez, J.M., Terol, J., Talon, M., Robles, M. 2005. Blast2GO: a universal tool for annotation, visualization and analysis in functional genomics research. Bioinformatics, 21(18), 3674–6.

Corteggiani Carpinelli, E., Telatin, A., Vitulo, N., Forcato, C., D’Angelo, M., Schiavon, R., Vezzi, A., Giacometti, G.M., Morosinotto, T., Valle, G. 2014. Chromosome scale genome assembly and transcriptome profiling of *Nannochloropsis gaditana* in nitrogen depletion. Mol Plant, 7(2), 323–35.

Deshpande, R.A., Shankar, V. 2002. Ribonucleases from T2 Family. Critical Reviews in Microbiology, 28(2), 79–122.

Doan, T.T.Y., Sivaloganathan, B., Obbard, J.P. 2011. Screening of marine microalgae for biodiesel feedstock. Biomass and Bioenergy, 35(7), 2534–2544.

Doerr, A. 2014. DIA mass spectrometry. Nature Methods, 12, 35.

Dray, S., Dufour, A.-B. 2007. The ade4 Package: Implementing the Duality Diagram for Ecologists. 2007, 22(4), 20.

Espie, G.S., Kimber, M.S. 2011. Carboxysomes: cyanobacterial RubisCO comes in small packages. Photosynth Res, 109(1-3), 7–20.

Fawley, M.W., Jameson, I., Fawley, K.P. 2015. The phylogeny of the genus *Nannochloropsis* (Monodopsidaceae, Eustigmatophyceae), with descriptions of *N. australis* sp. nov. and *Microchloropsis* gen. nov. Phycologia, 54(5), 545–552.

Fellenberg, K., Hauser, N.C., Brors, B., Neutzner, A., Hoheisel, J.D., Vingron, M. 2001. Correspondence analysis applied to microarray data. Proceedings of the National Academy of Sciences, 98(19), 10781–10786.

Flowers, T., Troke, P., Yeo, A. 1977. The mechanism of salt tolerance in halophytes. Annual review of plant physiology, 28(1), 89–121.

Flowers, T.J., Colmer, T.D. 2008. Salinity tolerance in halophytes. New Phytol, 179(4), 945–63.

Fulda, M., Schnurr, J., Abbadi, A., Heinz, E., Browse, J. 2004. Peroxisomal Acyl-CoA synthetase activity is essential for seedling development in *Arabidopsis thaliana*. Plant Cell, 16(2), 394–405.

Gao, W., Li, H.Y., Xiao, S., Chye, M.L. 2010. Acyl◻CoA◻binding protein 2 binds lysophospholipase 2 and lysoPC to promote tolerance to cadmium◻induced oxidative stress in transgenic *Arabidopsis*. The Plant Journal, 62(6), 989–1003.

Garg, A.K., Kim, J.-K., Owens, T.G., Ranwala, A.P., Do Choi, Y., Kochian, L.V., Wu, R.J. 2002. Trehalose accumulation in rice plants confers high tolerance levels to different abiotic stresses. Proceedings of the National Academy of Sciences, 99(25), 15898–15903.

Ge, X., Xia, Y. 2008. The role of AtNUDT7, a Nudix hydrolase, in the plant defense response. Plant signaling & behavior, 3(2), 119–120.

Georgianna, D.R., Mayfield, S.P. 2012. Exploiting diversity and synthetic biology for the production of algal biofuels. Nature, 488(7411), 329.

Gerhardt, B. 1992. Fatty acid degradation in plants. Prog Lipid Res, 31(4), 417–46.

Gilmour, D., Hipkins, M., Webber, A., Baker, N., Boney, A. 1985. The effect of ionic stress on photosynthesis in *Dunaliella tertiolecta*. Planta, 163(2), 250–256.

Goddijn, O.J., van Dun, K. 1999. Trehalose metabolism in plants. Trends in plant science, 4(8), 315–319.

Greenacre, M.J. 1984. Theory and applications of correspondence analysis.

Grieco, M., Tikkanen, M., Paakkarinen, V., Kangasjarvi, S., Aro, E.M. 2012. Steady-state phosphorylation of light-harvesting complex II proteins preserves photosystem I under fluctuating white light. Plant Physiol, 160(4), 1896–910.

Hershkovitz, N., Oren, A., Cohen, Y. 1991. Accumulation of trehalose and sucrose in cyanobacteria exposed to matric water stress. Appl. Environ. Microbiol., 57(3), 645–648.

Hibino, T., Waditee, R., Araki, E., Ishikawa, H., Aoki, K., Tanaka, Y., Takabe, T. 2002. Functional characterization of choline monooxygenase, an enzyme for betaine synthesis in plants. Journal of Biological Chemistry, 277(44), 41352–41360.

Hirth, M., Liverani, S., Mahlow, S., Bouget, F.-Y., Pohnert, G., Sasso, S. 2017. Metabolic profiling identifies trehalose as an abundant and diurnally fluctuating metabolite in the microalga *Ostreococcus tauri*. Metabolomics, 13(6), 68.

Ho, S.-H., Nakanishi, A., Kato, Y., Yamasaki, H., Chang, J.-S., Misawa, N., Hirose, Y., Minagawa, J., Hasunuma, T., Kondo, A. 2017. Dynamic metabolic profiling together with transcription analysis reveals salinity-induced starch-to-lipid biosynthesis in alga *Chlamydomonas* sp. JSC4. Scientific reports, 7, 45471.

Huertas, E., Montero, O., Lubián, L.M. 2000. Effects of dissolved inorganic carbon availability on growth, nutrient uptake and chlorophyll fluorescence of two species of marine microalgae. Aquacultural engineering, 22(3), 181–197.

Irani, S., Todd, C.D. 2016. Ureide metabolism under abiotic stress in *Arabidopsis thaliana*. J Plant Physiol, 199, 87–95.

Jambunathan, N., Mahalingam, R. 2006. Analysis of *Arabidopsis* growth factor gene 1 (GFG1) encoding a nudix hydrolase during oxidative signaling. Planta, 224(1), 1–11.

Jing, H., Takagi, J., Liu, J.-h., Lindgren, S., Zhang, R.-g., Joachimiak, A., Wang, J.-h., Springer, T.A. 2002. Archaeal surface layer proteins contain beta propeller, PKD, and beta helix domains and are related to metazoan cell surface proteins. Structure (London, England : 1993), 10(10), 1453–1464.

Juergens, M.T., Deshpande, R.R., Lucker, B.F., Park, J.-J., Wang, H., Gargouri, M., Holguin, F.O., Disbrow, B., Schaub, T., Skepper, J.N. 2015. The regulation of photosynthetic structure and function during nitrogen deprivation in *Chlamydomonas reinhardtii*. Plant physiology, 167(2), 558–573.

Karthikaichamy, A., Deore, P., Srivastava, S., Coppel, R., Bulach, D., Beardall, J., Noronha, S. 2018. Temporal acclimation of *Microchloropsis gaditana* CCMP526 in response to hypersalinity. Bioresource Technology, 254, 23–30.

Katz, A., Waridel, P., Shevchenko, A., Pick, U. 2007. Salt-induced changes in the plasma membrane proteome of the halotolerant alga *Dunaliella salina* as revealed by blue native gel electrophoresis and nano-LC-MS/MS analysis. Molecular & Cellular Proteomics, 6(9), 1459–1472.

Kav, N.N., Srivastava, S., Goonewardene, L., Blade, S.F. 2004. Proteome◻level changes in the roots of Pisum sativum in response to salinity. Annals of Applied Biology, 145(2), 217–230.

Kim, B.-H., Ramanan, R., Kang, Z., Cho, D.-H., Oh, H.-M., Kim, H.-S. 2016. *Chlorella sorokiniana* HS1, a novel freshwater green algal strain, grows and hyperaccumulates lipid droplets in seawater salinity. Biomass and Bioenergy, 85, 300–305.

Kim, Y.E., Hipp, M.S., Bracher, A., Hayer-Hartl, M., Hartl, F.U. 2013. Molecular chaperone functions in protein folding and proteostasis. Annu Rev Biochem, 82, 323–55.

Kishino, H., Waddell, P.J. 2000. Correspondence analysis of genes and tissue types and finding genetic links from microarray data. Genome Informatics, 11, 83–95.

Krishnaraj, S., Thorpe, T. 1996. Salinity stress effects on [14C-1]-and [14C-6]-glucose metabolism of a salt-tolerant and salt-susceptible variety of wheat. International journal of plant sciences, 157(1), 110–117.

Kumar, R., Subba, A., Kaur, C., Ariyadasa, T.U., Sharan, A., Pareek, A., Sopory, S.K., Singla-Pareek, S.L. 2018. OsCBSCBSPB4 is a Two Cystathionine-β-Synthase Domain-containing Protein from Rice that Functions in Abiotic Stress Tolerance. Curr Genomics, 19(1), 50–59.

Lai, C.Y., Cronan, J.E. 2004. Isolation and characterization of beta-ketoacyl-acyl carrier protein reductase (fabG) mutants of *Escherichia coli* and *Salmonella enterica* serovar Typhimurium. J Bacteriol, 186(6), 1869–78.

Lescano, C.I., Martini, C., Gonzalez, C.A., Desimone, M. 2016. Allantoin accumulation mediated by allantoinase downregulation and transport by Ureide Permease 5 confers salt stress tolerance to *Arabidopsis* plants. Plant Mol Biol, 91(4-5), 581–95.

Liska, A.J., Shevchenko, A., Pick, U., Katz, A. 2004. Enhanced Photosynthesis and Redox Energy Production Contribute to Salinity Tolerance in *Dunaliella* as Revealed by Homology-Based Proteomics. Plant Physiology, 136(1), 2806–2817.

Lu, C., Hills, M.J. 2002. *Arabidopsis* mutants deficient in diacylglycerol acyltransferase display increased sensitivity to abscisic acid, sugars, and osmotic stress during germination and seedling development. Plant Physiol, 129(3), 1352–8.

Martínez-Roldán, A., Perales-Vela, H.V., Cañizares-Villanueva, R.O., Torzillo, G. 2014. Physiological response of *Nannochloropsis* sp. to saline stress in laboratory batch cultures. Journal of applied phycology, 26(1), 115–121.

Mata-Pérez, C., Sánchez-Calvo, B., Padilla, M.N., Begara-Morales, J.C., Luque, F., Melguizo, M., Jiménez-Ruiz, J., Fierro-Risco, J., Peñas-Sanjuán, A., Valderrama, R. 2016. Nitro-fatty acids in plant signaling: nitro-linolenic acid induces the molecular chaperone network in *Arabidopsis*. Plant physiology, 170(2), 686–701.

McLean, M.D., Yevtushenko, D.P., Deschene, A., Van Cauwenberghe, O.R., Makhmoudova, A., Potter, J.W., Bown, A.W., Shelp, B.J. 2003. Overexpression of glutamate decarboxylase in transgenic tobacco plants confers resistance to the northern root-knot nematode. Molecular Breeding, 11(4), 277–285.

Mei, X., Chen, Y., Zhang, L., Fu, X., Wei, Q., Grierson, D., Zhou, Y., Huang, Y., Dong, F., Yang, Z. 2016. Dual mechanisms regulating glutamate decarboxylases and accumulation of gamma-aminobutyric acid in tea (*Camellia sinensis*) leaves exposed to multiple stresses. Sci Rep, 6, 23685.

Miernyk, J.A. 1999. Protein folding in the plant cell. Plant Physiol, 121(3), 695–703.

Mikami, K., Murata, N. 2003. Membrane fluidity and the perception of environmental signals in cyanobacteria and plants. Progress in lipid research, 42(6), 527–543.

Miller, G., Suzuki, N., Ciftci◻Yilmaz, S., Mittler, R. 2010. Reactive oxygen species homeostasis and signalling during drought and salinity stresses. Plant, cell & environment, 33(4), 453–467.

Naganuma, T., Nomura, N., Yao, M., Mochizuki, M., Uchiumi, T., Tanaka, I. 2010. Structural basis for translation factor recruitment to the eukaryotic/archaeal ribosomes. J Biol Chem, 285(7), 4747–56.

Naranjo, M.Á., Forment, J., Roldán, M., Serrano, R., Vicente, O. 2006. Overexpression of Arabidopsis thaliana LTL1, a salt-induced gene encoding a GDSL-motif lipase, increases salt tolerance in yeast and transgenic plants. Plant, Cell & Environment, 29(10), 1890–1900.

Ou, X., Ji, C., Han, X., Zhao, X., Li, X., Mao, Y., Wong, L.L., Bartlam, M., Rao, Z. 2006. Crystal structures of human glycerol 3-phosphate dehydrogenase 1 (GPD1). J Mol Biol, 357(3), 858–69.

Palleros, D.R., Reid, K.L., Shi, L., Welch, W.J., Fink, A.L. 1993. ATP-induced protein-Hsp70 complex dissociation requires K+ but not ATP hydrolysis. Nature, 365(6447), 664–6.

Pandit, P.R., Fulekar, M.H., Karuna, M.S.L. 2017. Effect of salinity stress on growth, lipid productivity, fatty acid composition, and biodiesel properties in *Acutodesmus obliquus* and Chlorella vulgaris. Environmental Science and Pollution Research, 24(15), 13437–13451.

Perrineau, M.M., Zelzion, E., Gross, J., Price, D.C., Boyd, J., Bhattacharya, D. 2014. Evolution of salt tolerance in a laboratory reared population of *Chlamydomonas reinhardtii*. Environmental microbiology, 16(6), 1755–1766.

Roberts, A., Gill, R., Hussey, R., Mikolajek, H., Erskine, P., Cooper, J., Wood, S., Chrystal, E., Shoolingin-Jordan, P. 2013. Insights into the mechanism of pyrrole polymerization catalysed by porphobilinogen deaminase: high-resolution X-ray studies of the *Arabidopsis thaliana* enzyme. Acta Crystallographica Section D: Biological Crystallography, 69(3), 471–485.

Schluepmann, H., Pellny, T., van Dijken, A., Smeekens, S., Paul, M. 2003. Trehalose 6-phosphate is indispensable for carbohydrate utilization and growth in *Arabidopsis thaliana*. Proceedings of the National Academy of Sciences, 100(11), 6849–6854.

Silveira, J.A.G., Carvalho, F.E.L. 2016. Proteomics, photosynthesis and salt resistance in crops: An integrative view. Journal of Proteomics, 143, 24–35.

Simionato, D., Sforza, E., Carpinelli, E.C., Bertucco, A., Giacometti, G.M., Morosinotto, T. 2011. Acclimation of Nannochloropsis gaditana to different illumination regimes: effects on lipids accumulation. Bioresource technology, 102(10), 6026–6032.

Sithtisarn, S., Yokthongwattana, K., Mahong, B., Roytrakul, S., Paemanee, A., Phaonakrop, N., Yokthongwattana, C. 2017. Comparative proteomic analysis of *Chlamydomonas reinhardtii* control and a salinity-tolerant strain revealed a differential protein expression pattern. Planta, 246(5), 843–856.

Subramanyam, R., Jolley, C., Thangaraj, B., Nellaepalli, S., Webber, A.N., Fromme, P. 2010. Structural and functional changes of PSI-LHCI supercomplexes of *Chlamydomonas reinhardtii* cells grown under high salt conditions. Planta, 231(4), 913–922.

Sugimoto, M., Takeda, K. 2009. Proteomic analysis of specific proteins in the root of salt-tolerant barley. Bioscience, biotechnology, and biochemistry, 73(12), 2762–2765.

Summer, E.J., Schmid, V.H., Bruns, B.U., Schmidt, G.W. 1997. Requirement for the H phosphoprotein in photosystem II of *Chlamydomonas reinhardtii*. Plant Physiol, 113(4), 1359–68.

Supek, F., Bošnjak, M., Škunca, N., Šmuc, T. 2011. REVIGO Summarizes and Visualizes Long Lists of Gene Ontology Terms. PLOS ONE, 6(7), e21800.

Suzuki, N., Bassil, E., Hamilton, J.S., Inupakutika, M.A., Zandalinas, S.I., Tripathy, D., Luo, Y., Dion, E., Fukui, G., Kumazaki, A., Nakano, R., Rivero, R.M., Verbeck, G.F., Azad, R.K., Blumwald, E., Mittler, R. 2016. ABA Is Required for Plant Acclimation to a Combination of Salt and Heat Stress. in: PloS one, Vol. 11, pp. e0147625.

Takabe, T., Incharoensakdi, A., Arakawa, K., Yokota, S. 1988. CO2 fixation rate and RuBisCO content increase in the halotolerant cyanobacterium, Aphanothece halophytica, grown in high salinities. Plant physiology, 88(4), 1120–1124.

Takagi, M., Yoshida, T. 2006. Effect of salt concentration on intracellular accumulation of lipids and triacylglyceride in marine microalgae *Dunaliella* cells. Journal of bioscience and bioengineering, 101(3), 223–226.

Troncoso-Ponce, M.A., Cao, X., Yang, Z., Ohlrogge, J.B. 2013. Lipid turnover during senescence. Plant Science, 205, 13–19.

Vizcaíno, J.A., Csordas, A., Del-Toro, N., Dianes, J.A., Griss, J., Lavidas, I., Mayer, G., Perez-Riverol, Y., Reisinger, F., Ternent, T. 2016. 2016 update of the PRIDE database and its related tools. Nucleic acids research, 44(D1), D447–D456.

Wada, H., Gombos, Z., Murata, N. 1994. Contribution of membrane lipids to the ability of the photosynthetic machinery to tolerate temperature stress. Proc Natl Acad Sci U S A, 91(10), 4273–7.

Wang, Y.P., Zhou, L.S., Zhao, Y.Z., Wang, S.W., Chen, L.L., Liu, L.X., Ling, Z.Q., Hu, F.J., Sun, Y.P., Zhang, J.Y., Yang, C., Yang, Y., Xiong, Y., Guan, K.L., Ye, D. 2014. Regulation of G6PD acetylation by SIRT2 and KAT9 modulates NADPH homeostasis and cell survival during oxidative stress. Embo j, 33(12), 1304–20.

Wei, S., Bian, Y., Zhao, Q., Chen, S., Mao, J., Song, C., Cheng, K., Xiao, Z., Zhang, C., Ma, W., Zou, H., Ye, M., Dai, S. 2017. Salinity-Induced Palmella Formation Mechanism in Halotolerant Algae *Dunaliella salina* Revealed by Quantitative Proteomics and Phosphoproteomics. Frontiers in Plant Science, 8(810).

Wijffels, R.H., Barbosa, M.J. 2010. An outlook on microalgal biofuels. Science, 329(5993), 796–799.

Wyn Jones, R. 1977. A hypothesis on cytoplasmic osmoregulation. Regulation of Cell Membrane Activities in Higher Plants., 121–136.

Young, J.N., Hopkinson, B.M. 2017. The potential for co-evolution of CO_2_-concentrating mechanisms and Rubisco in diatoms. Journal of experimental botany, 68(14), 3751–3762.

Zhu, B.-H., Zhang, R.-H., Lv, N.-N., Yang, G.-P., Wang, Y.-S., Pan, K.-H. 2018. The Role of Malic Enzyme on Promoting Total Lipid and Fatty Acid Production in *Phaeodactylum tricornutum*. Frontiers in Plant Science, 9(826).

